# ChemBERTaDDI: Transformer Driven Molecular Structures and Clinical Data for predicting drug-drug interactions

**DOI:** 10.1101/2025.01.22.634309

**Authors:** Anastasiya A. Gromova, Anthony S. Maida

## Abstract

The problem of polypharmacy arises when two or more drugs taken in combination cause adverse side effects, even when the use of the drugs individually causes no harm. Drug-drug interactions (DDIs) are a major cause of these reactions, contributing to increased morbidity and mortality. As the potential for harmful DDI grows combinatorially, the prediction of drug-drug interactions is increasingly critical for patient safety and effective healthcare management. In this paper, we present the *ChemBERTaDDI* framework a robust approach that uses transformer self-attention to extract latent molecular representations. By employing *ChemBERTa-77M-MLM*—a transformer-based language model pretrained on SMILES sequences—our approach generates enriched chemical embeddings that capture detailed molecular structural information. These embeddings are integrated with clinical mono side effect data and processed through a DNN predictor, enabling the learning of complex pairwise interaction patterns with minimal architectural overhead. Experiments performed using this combined data on a benchmark data set show superior performance compared with five state-of-the-art methods: Decagon, DeepDDI, MDF-SA-DDI, DPSP and NNPS. *ChemBERTaDDI* outperforms the baseline architectures, as measured by F1 and AUROC, and generalizes to new introduced drug compounds.

## I. Introduction

**P**OLYPHARMACY, the use of multiple drugs by patients, has become a widespread practice in modern healthcare. Although often medically necessary and offering therapeutic benefits, polypharmacy often leads to harmful drug-drug interactions (DDI). The challenge of predicting potential DDIs becomes increasingly complex as the number of possible drug combinations grows combinatorially, far outpacing the capacity of traditional clinical trials to identify rare and complex interactions. Existing computational methods for DDI prediction rely on extracting chemical and biological features from diverse drug properties. However, some drug properties are hard and costly to discover and may not be available in many cases. Therefore, reliable identification of DDIs is essential to mitigate adverse drug adverse events, particularly during the drug discovery and development process, which is important for both patient safety and pharmaceutical industry.

The landscape of DDI prediction methodologies spans a broad spectrum, ranging from traditional similarity-based approaches to deep learning techniques that use multiple data modalities. Traditional approaches in DDI prediction are built on the foundational principle that “similar drugs can interact with the same drug” [1]–[6]. These approaches extract drug characteristics such as chemical fingerprints, physical properties and phenotypic gene information, and then apply similarity-based measures for prediction. Recent advances in deep models have further transformed the field. Zitnick et al. [7] introduced Decagon, which transforms the DDI prediction task as a multi-relational link prediction problem on a multi-modal graph. Decagon uses a graph convolutional network (GCN) as an encoder and a tensor decomposition as decoder to learn robust node embeddings from interaction networks. Gomez-Bombarelli et al. [8] leveraged variational autoencoders (VAEs) to learn continuous and expressive representations of chemical structures, thereby enabling nuanced exploration of the chemical space. In parallel, natural language processing (NLP) techniques have emerged as a powerful tool for DDI prediction. Ishani Modal et al. [9] developed BERTChem-DDI, which formulates the DDI prediction problem as a relation classification task using transformer-based contextual embeddings. Ryu et al. [10] developed DeepDDI, which fuses chemical structure and drug names into a unified feature vector. The predictor is an eight-layer multilabel deep neural network (DNN) that predicts DDIs in a form of human-readable DDI prediction sentences. Huang et al. [11] introduced Caster, which integrates multi-source drug fusion with sequential pattern mining to uncover latent drug interactions, while Han et al. [12] designed SmileGNN, an architecture that combines drug structural information with topological features extracted from a knowledge graph via a graph neural network (GNN). Li et al. [13] developed DSNDDI, a dual-view framework that integrates local drug substructure information with global drug pair interactions, the resulting embeddings are fed into DSN decoder for DDI prediction. Kim et al. [14] proposed DeSIDE-DDI, which uses a drug-induced gene expression profile along with compound structural and property data for interpretable DDI prediction. Finally, Feng et al. [15] introduced DPDDI, which uses a graph convolution network (GCN) to capture low-dimensional topological relationships of drugs with a deep neural network (DNN) model as predictor. Despite these advances, many methods continue to depend heavily on handcrafted features and suffer from challenges related to noise in heterogeneous graphs, thus limiting prediction accuracy and scalability. Addressing these issues remains a critical research direction in DDI prediction systems.

To address the above limitations, this work focuses on the *ChemBERTaDDI* framework, which improves drug-drug interaction (DDI) prediction by integrating transformer-based language model embeddings with clinical mono side effect data. Our approach leverages domain specific contextual embeddings *ChemBERTa-10M-MLM* [16] and *ChemBERTa-77M-MLM* embeddings [17] derived from transformer attention mechanisms based on foundational BERT architecture [18] and its robust adaptation, RoBERTa [19]. These context-full representations are then concatenated with individual drug clinical side effect features to form a comprehensive multi-modal drug representation. The predictor leverages combined features derived from the molecule’s latent continuous vector representation—capturing its chemical properties—and its mono side effect vector. This comprehensive feature set is then fed into a multilayer neural network for binary classification. By maintaining frozen-layer embeddings, our *ChemBERTaDDI* framework efficiently captures high-quality molecular representations without extra computational overhead. The framework predicts whether a pair of drugs will result in specific adverse side effects.

Benchmark comparisons with DeepWalk, Decagon, and NNPS demonstrate that *ChemBERTaDDI* outperforms existing models in DDI prediction. Additionally, a single neural network model is trained across 964 polypharmacy side effects, utilizing a comprehensive 52*K* token SMILES [20] vocabulary to ensure scalability and adaptability to new drug compounds. In practice, our *ChemBERTaDDI* framework uses a single neural network model trained on 964 polypharmacy side effects while preserving ChemBERTa’s pre-trained knowledge, thereby reducing computational overhead. This approach allows the framework to focus on learned molecular representations within a comprehensive 52*K* token SMILES chemical vocabulary, ensuring adaptability to new compound data and scalability. The complete source code and dataset used in this study are available on GitHub (https://github.com/anastasiyagromova/ChemBERTaDDI) for reproducibility purposes. Key contributions of this paper are summarized as follows:

- Developed the *ChemBERTaDDI* framework (Section 2) that leverages drug clinical data along with external chemical structure embeddings (Section 3) for DDI prediction.
- Demonstrated that incorporating SMILES-derived molecular insights with drug mono side effect data significantly improves DDI prediction performance (Section 5). Moreover, our ablation studies comparing *ChemBERTa-10M-MLM* [16] and *ChemBERTa-77M-MLM* [17] models further reveal that model scale plays a critical role in enhancing predictive accuracy (Section 6).

## II.Methodology

Given a set of 645 drugs, denoted by *D* = {*d*_1_, *d*_2_, …, *d*_645_} and 964 polypharmacy side effects, denoted by *S* = {*s*_1_, *s*_2_, …, *s*_964_}, our objective is to determine whether an interaction occurs for a pair of drugs (*d*_*i*_, *d*_*j*_) and to accurately predict the specific side effect(s) if present.

For each side effect *s* ∈ *S*, we construct a symmetric adjacency matrix *A* ∈ ℝ^*m×m*^, where *m* = 645 (the total number of drugs). Each entry *a*_*ij*_ is defined as:

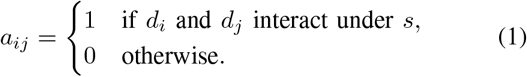

Our primary objective is to develop a robust predictive model that leverages both high-dimensional clinical data with generalizable rich chemical structure representations to accurately predict DDIs.

### A. ChemBERTaDDI Framework

The **ChemBERTaDDI** framework integrates clinical mono side effect features with transformer-based molecular embeddings to predict potential DDIs. The framework operates in the following steps. Refer to Figure 2.

**Fig. 1.**
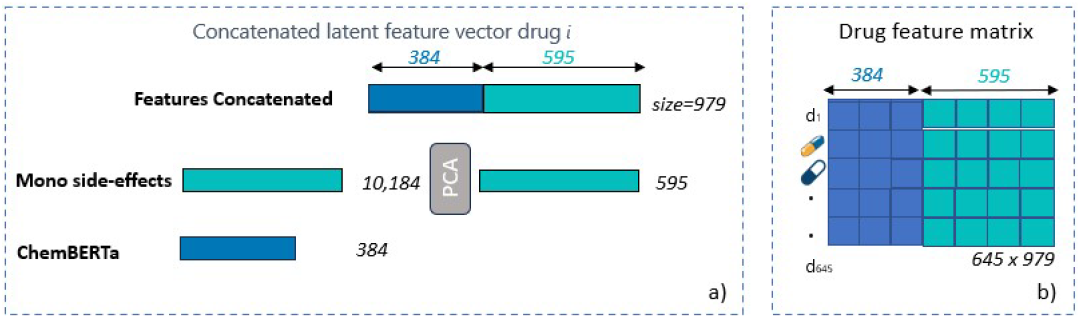
ChemBERTaDDI Feature Representations. (a) For each drug *d*_*i*_, the feature vector is a concatenation of two modalities: the 384-dimensional ChemBERTa drug–chemical structure embedding (blue) and the 595-dimensional PCA-transformed representation of mono side effects (green). Each cell in the feature matrix encapsulates this integrated multimodal representation. (b) The complete drug feature matrix is constructed by stacking these concatenated vectors for all 645 drugs, resulting in a 645 *×* 979 matrix. Each row represents a drug’s combined chemical and clinical feature vector, enabling downstream drug interaction prediction.

**Fig. 2.**
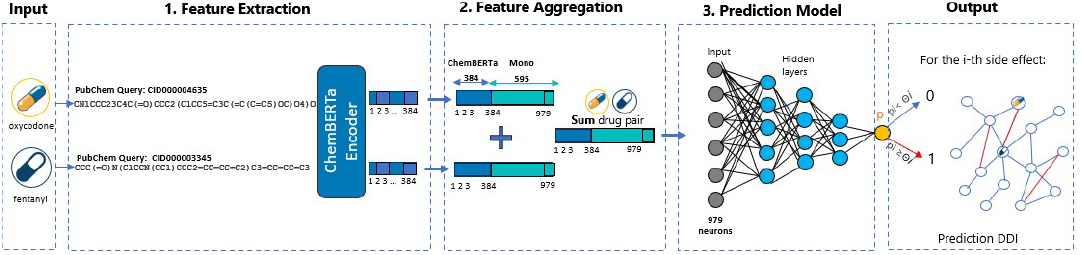
ChemBERTaDDI Framework. Step 1: Extract 645 unique drugs and retrieve their corresponding SMILES strings from PubChem. Transform each SMILES string into 384-dimensional feature embedding using ChemBERTa. Step 2: Aggregate features by concatenating the ChemBERTa embeddings (blue) with PCA-reduced mono side-effect vectors (green). For each drug pair, sum the two 979-dimensional vectors into a single comprehensive representation, which is then fed into a prediction model that outputs a sigmoid probability.

Building on the motivations and problem settings identified by Zitnick et al. (Decagon) [7] and Masumshah et al. (NNPS) [21], the objective is to predict drug-drug interactions (DDIs) associated with 964 side effects. Two key drug-related characteristics are explored (see Figure 1 Section (a) for details):

#### 1) Drug mono-side effects clinical information

Clinical information is represented as categorical data with a total of 10,184 categories. One-hot encoding is used to transform categorical variables into a binary vector, where each unique category is represented by a distinct binary feature to preserve the categorical nature. Principal component analysis (PCA) is applied for dimensionality reduction. While retaining 99% of the variance, PCA projects the data into a lower-dimensional space with 595 dimensions, effectively capturing the most significant patterns while greatly reducing computational complexity.

#### 2) Chemical structure information

Motivated by Huang et al. (CASTER) [11], which models drug pairs as a collection of chemical functional sub-structures to predict DDIs, we utilize chemical feature representations. The *ChemBERTaDDI* framework leverages the Simplified Molecular Input Line Entry System (SMILES) strings to capture the intricate chemical structures of drugs. SMILES serves as a textual notation for chemical compounds, encapsulating their molecular composition and structural configurations. SMILES entries are collected from the external PubChem database [22] and are tokenized using `AutoTokenizer` from Hugging Face’s Transformers library. `AutoTokenizer` is the sub-word tokenization technique that splits SMILES strings into tokens, allowing the transformer model to process them. Below is the summary of the *ChemBERTaDDI* components, refer to Figure 2 for details.

- **Feature Extraction:** The `AutoTokenizer` tokenizer converts SMILES strings into token IDs, which are then fed into the *ChemBERTa-77M-MLM* transformer model developed by Chithranda et al. [17]. The idea is similar to CASTER [11], instead using a frequent sequential pattern mining algorithm, we leverage `AutoTokenizer`, which generates the recurring chemical sub-structures of molecular representations of drugs. Inspired by the advancements in self-supervised pretraining on unlabeled SMILES data via a masked language modeling (MLM) for molecular property prediction [17], *ChemBERTaDDI* uses *ChemBERTa-77M-MLM* embeddings generated via transformer attention mechanisms. The *ChemBERTa* model processes tokens through multiple encoder layers to generate contextualized molecular embeddings. Serving as a robust feature extractor, *ChemBERTa-77M-MLM* transforms SMILES inputs into generalizable embeddings that effectively capture the chemical structure and properties of each drug based on its molecular composition.
- **Feature Aggregation:** The PCA-reduced clinical mono side effect features are concatenated with *ChemBERTa* embeddings to form a combined 979-dimensional feature vector for each drug. Subsequently, for each drug pair, their respective individual feature vectors are aggregated using an element-wise summation operation. The summation combines the attributes of both drugs, resulting in a feature vector that captures the joint clinical and chemical characteristics necessary for predicting DDIs.
- **Prediction Model:** Motivated by the neural network-based predictor (NNPS) introduced by Masumshah et al. [21], we aggregate multi-modal feature vectors that capture both clinical and chemical information and feed them into a single deep neural network for binary classification. This unified model is trained to predict the likelihood of adverse DDIs by leveraging the integrated feature representation, thereby maximizing predictive accuracy.

## III. Drug Feature Representations

Each drug *d*_*k*_ is represented by two complementary feature matrices, encoding both clinical information (*mono-side effects*) and chemical structure (via *ChemBERTa*). Specifically, for each side effect, two types of feature matrices are considered, refer to Figure 1: (1) the drug mono side effects matrix 645×10,184 (2) the drug chemical feature matrix 645×384 derived from ChemBERTa.

### A) Mono-side Effect Features

The individual drug mono side effects clinical information is represented via a one-hot encoded vector of length 10,184, indicating the presence or absence of specific mono side effects. Because the mono-side effect feature matrix is high-dimensional and sparse, the Principal Component Analysis (PCA) is applied to reduce dimensionality while retaining 99% of the variance. As a result, the reduced mono side effect vector yields:

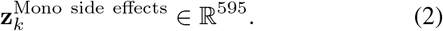

### B) Molecular Feature Extraction

For the purpose of modeling detailed molecular semantics of each drug, the proposed idea is to adapt *ChemBERTa-77M-MLM*, a transformer-based language model pretrained on unlabeled SMILES data. The decision to use *ChemBERTa-77M-MLM* over latent variable models stems from its ability to provide robust and contextually rich embeddings with fewer architectural complexities. Transformer models like *ChemBERTa* integrate self-attention mechanisms within a single, unified architecture—removing the need for separate encoder–decoder pipelines and complex reconstruction objectives common in autoencoders and variational frameworks. This streamlined design not only reduces the architectural burden but also generates robust, contextually enriched embeddings directly from unlabeled SMILES data, enhancing molecular semantic representation with fewer computational overheads and tuning challenges. By using transformer-based models, *ChemBERTa* offers a powerful tool for capturing the nuanced semantics of molecular data, facilitating more accurate and efficient predictions in the drug-drug interaction (DDI) task. In the *ChemBERTaDDI* framework, *ChemBERTa-77M-MLM* acts as an encoder from which a continuous latent representation is obtained, where molecular representation of drugs has been used as both input and output, thus learning a 52,000-token vocabulary of molecular fragments. Chithrananda et al. [17] converted the discrete SMILES representations of the drug molecules into continuous multi-dimensional representation using *ChemBERTa-77M-MLM* transformer model. The primary objective of *ChemBERTa-77M-MLM* is to learn meaningful and robust representations of molecular structures encoded as SMILES strings. This is achieved through the Masked Language Modeling (MLM) paradigm, a foundational training objective borrowed from natural language processing models, such as BERT [18] and RoBERTa [19]. The *ChemBERTa-77M-MLM* model [17], employs the MLM paradigm, where 15% of the SMILES tokens in the input SMILES string are randomly masked/hidden, and the model is trained to predict these masked tokens based on their surrounding context. Formally, given a SMILES string represented as a sequence of tokens:

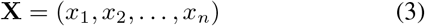

a subset of tokens is masked to produce:

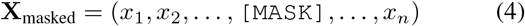

The objective is to maximize the likelihood of the masked tokens given the unmasked context:

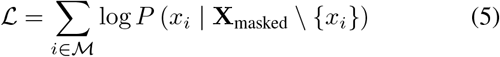

where:

- ℳ denotes the set of masked token indices.
- *P* (*x*_*i*_| ·) represents the probability of token *x*_*i*_ being the correct token given the surrounding context.

Let *f* (·) : *X* → *H* denote ChemBERTa’s encoder mapping a tokenized SMILES *X* (length ≤ 512) to a final hidden state *H* ∈ ℝ^384^. Using a *masked language modeling* objective ChemBERTa learns chemically meaningful latent representations from large-scale SMILES corpora. In our framework, we freeze ChemBERTa trained weights and extract the hidden representation molecular knowledge for each drug represented by a 384-dimensional embedding vector:

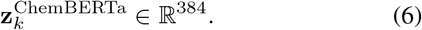

### C) Drug Pair Feature Representation

To facilitate DDI prediction, we concatenate the reduced mono-side effect features and the ChemBERTa embeddings for each drug, resulting in a following unified feature vector:

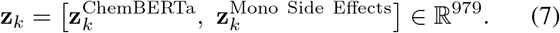

For a drug pair (*d*_*i*_, *d*_*j*_), we aggregate their respective feature vectors by summation:

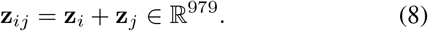

This aggregated feature vector serves as input to the neural network model, enabling the prediction of potential DDIs based on the combined clinical and chemical characteristics of the drug pair. However, it is important to note that such a summation may result in some information loss, a trade-off that warrants further investigation.

## IV. Prediction Model

In the *ChemBERTaDDI* framework, drug-drug interactions (DDIs) are predicted using a three hidden layer feedforward neural network for binary classification. The model processes an aggregated feature vector **z**_*ij*_ ∈ ℝ^979^ that combines chemical and clinical attributes. It consists of three fully connected hidden layers with 250, 300, and 200 neurons respectively, with weights initialized with the Glorot normal initializer and followed by Batch Normalization, ReLU activation, and Dropout regularization. The output layer contains a single neuron with a sigmoid activation function, producing a binary classification output indicating the presence (1) or absence (0) of an adverse DDI.

Let *F* (·) denote our NN-based predictor. For side effect *s*, the predicted interaction probability between (*d*_*i*_, *d*_*j*_) is:

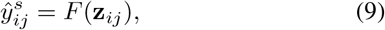

where 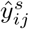 is compared against a threshold Θ to decide whether *s* occurs.

## V. Experimental Setup

This section explains the datasets and the experiments conducted using *ChemBERTaDDI*. Specifically, we integrated drug pair interactions (63, 473 pairs) and associated polypharmacy side effects, individual drug mono side effects, drug-protein interaction data, and SMILES representations for 645 drugs. Table I presents the statistical overview of the databases, while Table II summarizes the pre-processed feature dataset. The experiments aim to evaluate the performance of *ChemBERTaDDI* by addressing the following two questions: **Q1)** Do aggregated transformer embeddings achieve more accurate predictions of drug-drug interactions compared to other strong baseline models? **Q2)** Can aggregated transformer embeddings provide accurate DDI predictions using only drug chemical structure embeddings from *ChemBERTa*, without incorporating drug clinical characteristics?

**TABLE I.** Drug and Side Effect Databases.

| Database | Details |
| --- | --- |
| **TWOSIDES** | No. of side effects = 1,317<br>No. of drugs = 645<br>No. of drug pairs = 63,473 |
| **SIDER, OFF-SIDES** | No. of SIDER drugs = 1,556<br>No. of SIDER side effects = 5868<br>No. of OFFSIDES drugs = 1,332<br>No. of OFFSIDES side effects = 10,097 |
| **STITCH** | No. of compounds = 519,022<br>No. of protein targets = 8,934<br>No. of interactions = 8,083,600 |
| **PubChem** | No. of SMILES = 645 |

**TABLE II.** DB1 Dataset details.

| Dataset | Details | Features |
| --- | --- | --- |
| **DBI** | No. of polypharmacy effects = 964<br>No. of drugs = 645<br>No. of interactions = 63,473<br>No. of mono side effects = 10,184<br>No. of SMILES = 645 | No. PCA mono side effects = 595<br>No. ChemBERTa = 384<br>No. Drug feature = 979 |

### A. Databases

The annotated drug pairs with polypharmacy side effects are sourced from the TWOSIDES database [23], a comprehensive database with drug-drug interactions (DDIs) containing 1, 317 polypharmacy side effects across 645 drugs and 63, 473 drug pairs extracted from the FDA Adverse Event Reporting System (FAERS). In total the TWOSIDES database contains 4, 576, 785 associations, which include multiple instances of interactions between the same pairs of drugs. Next, OFFSIDES and SIDER databases are integrated to derive the side effects of individual drugs (mono side effects). The SIDER database [24] compiles side effects from drug labels, while the OFFSIDES database [23] includes off-label side effects observed during clinical trials. The SIDER information is extracted from drug labels and includes 1, 556 drugs and 5, 868 side effects compiled from public documents. The OFFSIDES information is collected from clinical trials and includes 1, 332 drugs and 10, 097 off-label side effects. By merging these databases and eliminating synonym side effects, 10, 184 mono side effects are obtained for the 645 drugs which are in TWOSIDES database. In addition, during the pre-processing, the chemical structure information for 645 drugs is obtained from the *PubChem* database [22], the largest publicly available database for chemical structure information. The Simplified Molecular Input Line Entry System (SMILES) [20] notation is used to represent the chemical structures of the 645 drugs in a string format. PubChem [22] is used to obtain the corresponding SMILES notation along with the drug names, which encodes molecular structure information as a string.

### B. DB1 Dataset

The above databases were adapted and pre-processed to construct comprehensive *DB1* dataset by preprocessing multiple sources to combine chemical and clinical information for DDI prediction. The *DB1* dataset is structured based on TWOSIDES annotated DDIs consisting of Polypharmacy Side Effects and a set of Drug Pairs. The polypharamacy side effects filtered to the 964 most frequently occurring side effects, which occurred in at least 500 DDIs. There are 63, 473 unique drug pairs; however since each drug pair can be associated with multiple adverse effects, the total number of training associations reaches 4, 576, 785. In summary, DB1 is structured as triplets (drug *i*, drug *j*, polypharmacy side effect *r*). In addition, we supplement *DB1* with mappings: 645 drugs to their corresponding PubChem SMILES string and drug names [22], and further associate each drug with a set of 10, 184 mono side effects. Further detailed information on the *DB1* dataset and appropriate features is available in Table II. Following standard practice outlined in [7], [21], known drugdrug interaction pairs from TWOSIDES are used as positive samples to build the positive set, while all the remaining unlabeled drug-drug pairs are considered as negative samples.

### C. Training Procedure

In the *ChemBERTaDDI* framework, pre-trained contextual embeddings *ChemBERTa10M-MLM* [16] and *ChemBERTa-77M-MLM* [17] are used to initialize the chemical representations of drug structures. These representations reside in a latent space, enabling generalization to novel drugs. For our training procedure, the selected compute was 64 cores, 440 GB RAM, 2816 GB disk space. We train a neural predictor via supervised learning on labeled drug pairs, using their chemical representations and mono side-effect information. Binary cross-entropy loss quantifies errors, optimized by Adam. Dropout and early stopping (10 epochs without validation improvement) mitigate overfitting. We measure performance (ROC, AUPR, Recall, Precision, F1-Score) for 964 side effects. The model is a feed-forward network with three hidden layers and a single output neuron. Its input layer has 979 neurons (a 384-dimensional *ChemBERTa* embedding plus 595-dimensional mono side effects). We split drug pairs by side effect into training (90%), validation (5%), and test (5%) sets, following a 5-fold crossvalidation. We explored three- and five-layer architectures, using both *ChemBERTa-10M-MLM* [16] and *ChemBERTa-77M-MLM* [17], with learning rates (0.001, 0.005, 0.01) and dropout rates (0.1, 0.2, 0.3). The best configuration uses a batch size of 1024, 50 epochs, dropout 0.3, and early stopping at 10 stalled epochs. Our final model (three fully connected layers of 250, 300, and 200 neurons) with *(ChemBERTa-77M-MLM, Mono)* features achieves an F1-score of 0.938, AUPR of 0.954, and AUROC of 0.965.

## VI. Results and Discussion

In this section, we present a comprehensive evaluation of the proposed *ChemBERTaDDI* framework, which combines clinical side effects with transformer-based chemical structure embeddings for drug-drug interaction (DDI) prediction. *ChemBERTaDDI* framework uses clinical side effect features along with transformer-based chemical structure embeddings from *ChemBERTa-77M-MLM* (pretrained on 77 million unique SMILES). By integrating complementary chemical and clinical insights, *ChemBERTaDDI* harnesses rich drug information to robustly predict drug-drug interactions.

Our approach was benchmarked against five previous methods: Decagon [7], NNPS [21], DeepDDI [10], MDF-SA-DDI [25], and DPSP [6]. As detailed in Table III, *ChemBERTaDDI* achieves a superior F-score of 0.937, an AUPR of 0.954, and AUROC of 0.967-substantially outperforming competing methods that range from F score of 0.727 to 0.936. Particularly, our framework not only improves predictive accuracy but also demonstrates robust performance in distinguishing interactions with dangerous polypharmacy side effects, as highlighted in the case study in Table IV.

**TABLE III.** Baseline Model Performance Comparison.

| Method | AUROC | AUPR | Precision | F1 Score |
| --- | --- | --- | --- | --- |
| ChemBERTa-77M-MLM, Mono | 0.965 | 0.954 | 0.909 | **0.937** |
| ChemBERTa-77M-MLM, Mono, Protein | 0.965 | 0.954 | 0.909 | **0.937** |
| Decagon | 0.874 | 0.825 | 0.771 | 0.850 |
| NNPS | 0.966 | 0.953 | 0.901 | 0.936 |
| DeepDDI | 0.996 | 0.889 | 0.727 | 0.727 |
| MDF-SA-DDI | 0.999 | 0.965 | 0.883 | 0.883 |
| DPSP | 0.999 | 0.9773 | 0.930 | 0.930 |

**TABLE IV.** Case Study: Results of Dangerous Polypharmacy Side Effects in ChemBERTaDDI and Decagon on AUPR.

| Polypharmacy | AUPR (ChEMBERTaDDI) | AUPR (Decagon) | Evidence |
| --- | --- | --- | --- |
| Sarcoma | 0.978 | 0.789 | Serban et al. [26] |
| Carcinoma of the Cervix | 0.952 | 0.810 | Arbyn et al. [27] |
| Malignant Hypertension | 1.00 | 0.858 | Januszewicz et al. [28] |
| Oophorectomy | 0.974 | 0.911 | Evans et al. [29] |
| Leukopenia | 0.960 | 0.695 | Flanagan et al. [30] |

To further probe the model (**Q2)**), we conducted ablation studies to assess the contribution of individual feature sets. Our results in Table V reveal that while the combination of *ChemBERTa* embeddings with mono side-effect features offers the best performance, the addition of protein data does not yield any measurable advantage. Moreover, when evaluated in isolation, the *ChemBERTa-77M-MLM* embeddings (trained on 77 million unique SMILES) achieve an average F1-score of 0.891, underscoring the necessity of integrating clinical information to achieve better performance. From a practical standpoint, an overall F1 score of approximately 0.94 is highly competitive, indicating a strong balance between precision and recall. However, to strengthen our conclusions and increase the clinical impact of our work, we further analyzed:

- Examining the worst-performing relationship types (as detailed in Table VII) to identify potential subgroups of side effects with ambiguous or sparse representations; such an analysis could drive improvements in data augmentation or hierarchical modeling.

**TABLE V.** ChemBERTaDDI Performance with various Feature Sets.

| Method | ROC | AUPR | Recall | Precision | F1 Score |
| --- | --- | --- | --- | --- | --- |
| ChemBERTa10M-MLM | 0.9281 | 0.9106 | 0.9375 | 0.8485 | 0.8887 |
| ChemBERTa10M-MLM, Protein | 0.9311 | 0.9144 | 0.9387 | 0.8530 | 0.8917 |
| ChemBERTa10M-MLM, Mono | 0.9630 | 0.9499 | 0.9680 | 0.9032 | 0.9336 |
| ChemBERTa10M-MLM, Mono, Protein | 0.9626 | 0.9491 | 0.9665 | 0.9048 | **0.9338** |
| ChemBERTa77M-MLM | 0.9308 | 0.9137 | 0.9377 | 0.8513 | 0.8909 |
| ChemBERTa77M-MLM, Protein | 0.9414 | 0.9272 | 0.9402 | 0.8685 | 0.9012 |
| ChemBERTa77M-MLM, Mono | 0.9653 | 0.9539 | 0.9681 | 0.9089 | **0.9368** |
| ChemBERTa77M-MLM, Mono, Protein | 0.9658 | 0.9538 | 0.9688 | 0.9081 | 0.9366 |

**TABLE VI.** Probing the Chemical Structure Embeddings.

| Models | Method | Average F1 |
| --- | --- | --- |
| **ChemBERTa-10M-MLM** | ChemBERTaDDI | 0.8887 |
| **ChemBERTa-77M-MLM** | ChemBERTaDDI | **0.8909** |

**TABLE VII.** The average of the best results for ChemBERTaDDI, NNPS, and Decagon methods for 964 side effects.

| Method | TP | FP | TN | FN | Precision | Recall | F-score |
| --- | --- | --- | --- | --- | --- | --- | --- |
| NNPS | 105,153 | 12,013 | 95,955 | 2,608 | 0.901 | **0.976** | 0.936 |
| Decagon | 208,963 | 54,855 | 163,988 | 9,880 | 0.771 | 0.950 | 0.850 |
| ChEMBERTaDDI | 107,790 | 12,817 | 99,061 | 4,116 | **0.909** | 0.970 | **0.938** |

**TABLE VIII.** Best Performing Polypharmacy Side Effects.

| Side Effect Name | AUROC | F Score | Loss |
| --- | --- | --- | --- |
| **Breast Enlargement** | 1.0 | 1.0 | 0.021 |
| **Mumps** | 1.0 | 1.0 | 0.014 |
| **Varicose Vein** | 1.0 | 1.0 | 0.035 |
| **Parotid Gland Enlargement** | 1.0 | 1.0 | 0.184 |
| **Epidural Hematoma** | 1.0 | 1.0 | 0.175 |

**TABLE IX.** Worst Performing Polypharmacy Side Effects.

| Side Effect Name | AUROC | F Score | Loss |
| --- | --- | --- | --- |
| **Polyarthrititis** | 0.924 | 0.880 | 0.419 |
| **Cholecystitis Acute** | 0.934 | 0.901 | 0.345 |
| **Diabetic Neuropathy** | 0.940 | 0.941 | 0.241 |
| **Primary Biliary Cirrhosis** | 0.944 | 0.966 | 0.203 |
| **Lung Abscess** | 0.957 | 0.960 | 0.216 |

In summary, while an F1 score of 0.938 is an impressive achievement, our detailed additional analyses, particularly on calibration, error patterns, and novel DDI discovery, underscore the nuanced strengths—and potential clinical utility—of *ChemBERTaDDI*. These further explorations not only boost confidence in our model’s predictive performance but also open new avenues for research in drug safety.

## VII. Conclusion

This paper developed a new computational *ChemBERTaDDI* framework for drug-drug interaction prediction that effectively integrates high-dimensional clinical data with transformer-based chemical molecular embeddings. The *ChemBERTaDDI* framework leverages a SMILES-based *ChemBERTa* encoder to extract a 384-dimensional embedding, which is then concatenated with a 595-dimensional mono side-effect feature vector before being passed into a feed-forward neural network. Experiments reveal two main findings: (i) combining molecular structure information from SMILES with drug mono side-effect data substantially improves DDI prediction performance; and (ii) relying solely on chemical embeddings is insufficient. By achieving state-of-the-art results on benchmark dataset, *ChemBERTaDDI* demonstrates strong generalization capability in scenarios where both chemical and clinical features are available. Future work will focus on incorporating additional modalities such as genetic or proteomic data, and exploring deeper transformer model to further enrich predictive power and scalability for DDI task.

## A. Anastasiya A. Gromova

Anastasiya Gromova received her Bachelor of Science in Engineering in 2015 and Master’s degree in Computer Engineering in 2017 from the University of Louisiana at Lafayette. She is currently a Ph.D. candidate in Computer Engineering at the same university. Her research interests include application of machine learning, artificial intelligence, natural language processing (NLP), computer vision, and deep learning techniques to advance medical decision-making in healthcare and bioinformatics.

**Fig. 3.**
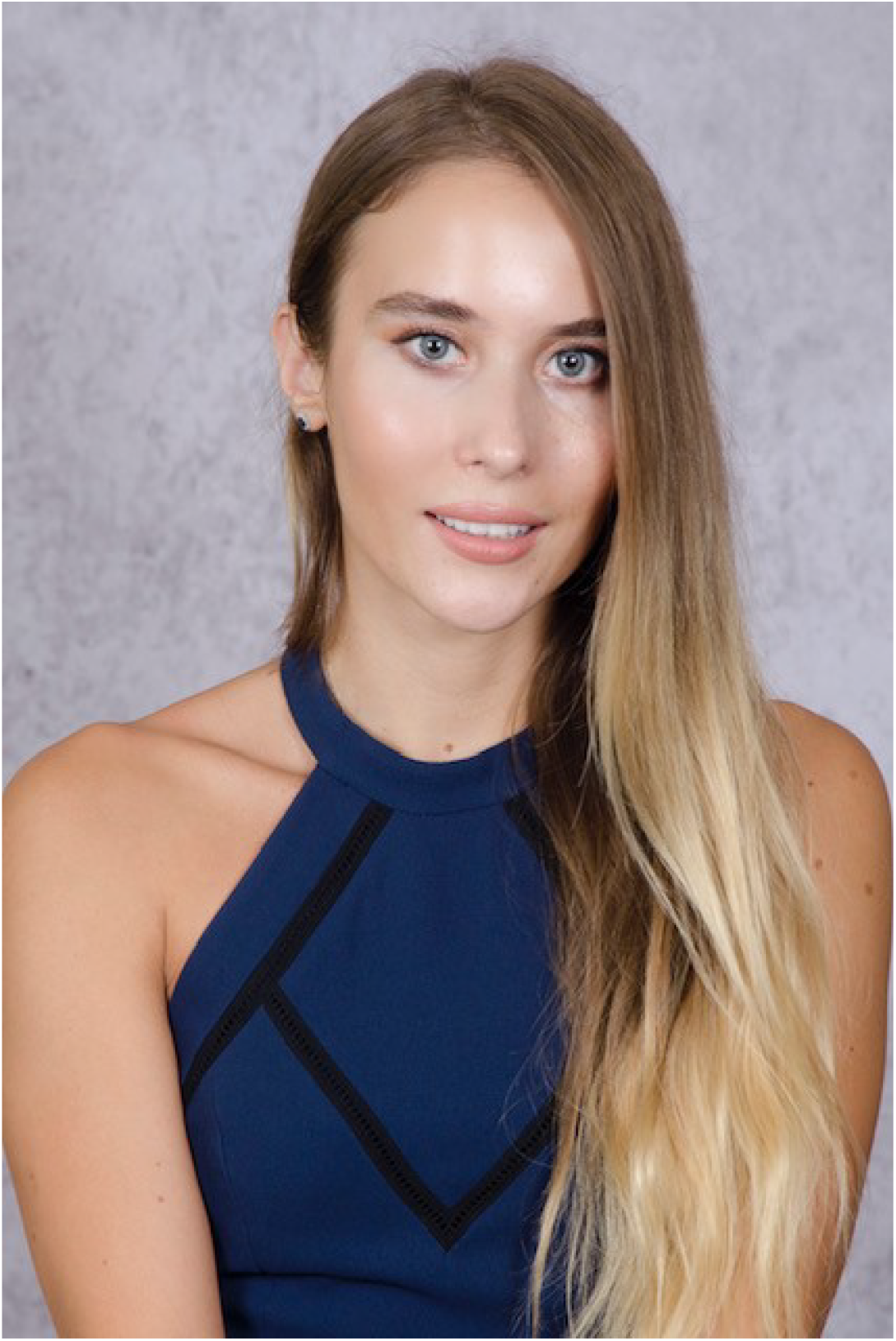
Anastasiya Gromova.

## B. Anthony S. Maida

Anthony S. Maida received the BA in mathematics, MS in computer science, and PhD in psychology, all from the State University of New York at Buffalo. He received postdoctoral training at Brown University and the University of California at Berkeley. He taught a the Pennsylvania State University and is currently an Associate Professor at the School of Computing and Informatics, the University of Louisiana at Lafayette. His research interests are in biologically inspired artificial intelligence, brain simulation, and deep learning.

**Fig. 4.**
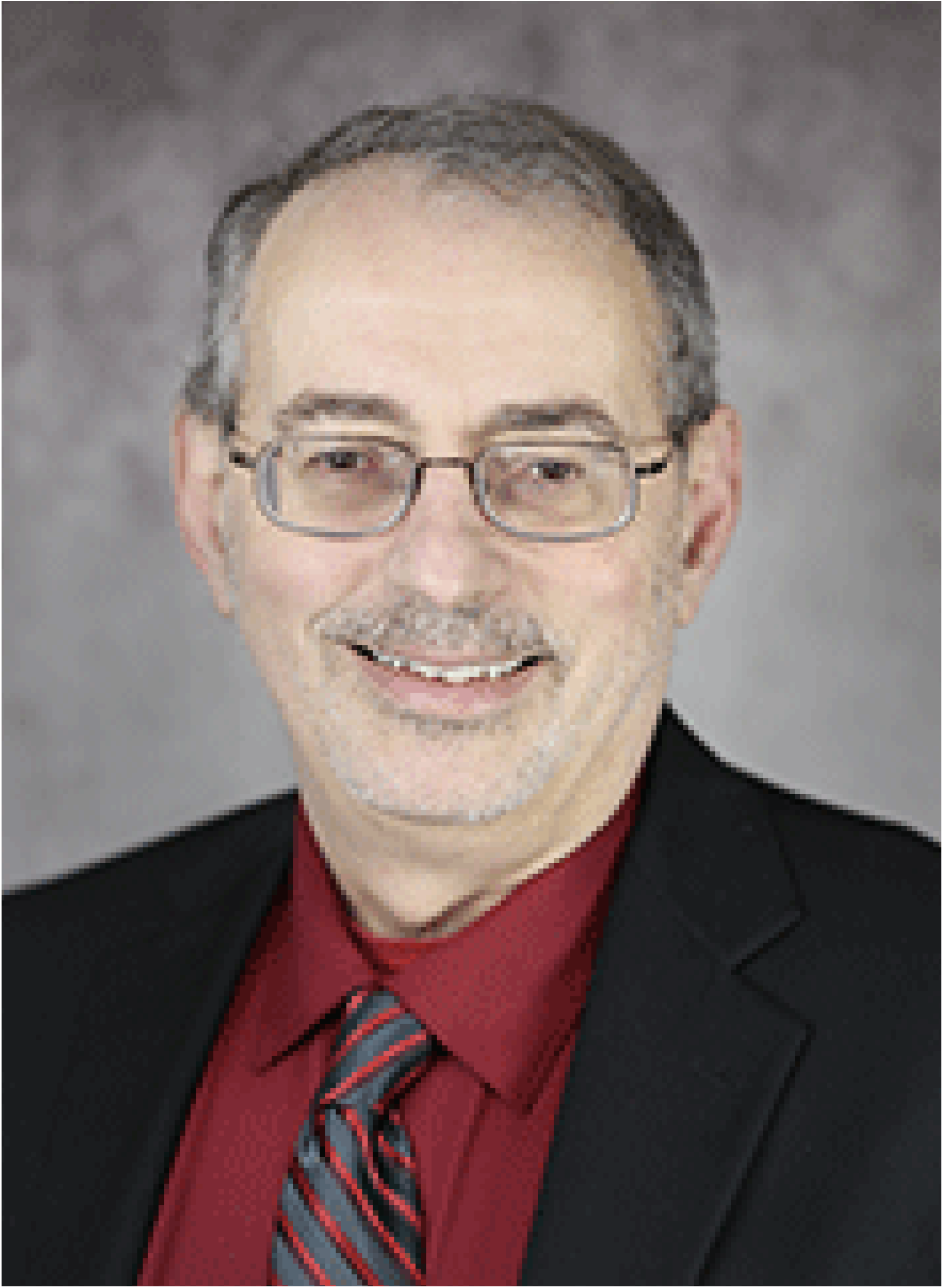
Anthony S. Maida.

